# The aging transcriptome: a condition-dependent response to a deteriorating soma

**DOI:** 10.64898/2025.12.22.696078

**Authors:** Zeeshan A. Syed, Martin I. Brengdahl, Vinesh N. Shenoi, Avani Mital, Christopher M. Kimber, Helen Ekman, Claudio Mirabello, Malin Larsson, David Berger, Urban Friberg

## Abstract

Organismal function requires precise gene expression, as deviations reduce fitness and can cause disease. The widespread expression changes characteristic of old age therefore suggests that aging itself may partly stem from gene dysregulation. Alternatively, many of these changes may represent a plastic response to somatic decline, tuning the organism to an altered physiological state. We tested the latter hypothesis by experimentally reducing the condition of *Drosophila melanogaster* females independently of age and comparing the resulting expression changes with those occurring naturally during aging. Consistent with the plasticity hypothesis, we find substantial overlap between genes responding to reduced condition and old age. Downregulated genes are enriched for metabolic functions, with a consistent, albeit weaker, association with mitochondrial function and cytoplasmic translation, while upregulated genes relate to genome maintenance. Both old age and reduced condition also cause downregulation of tissue-specific and female-biased genes, as expected when energy is reallocated to core cellular processes. In line with a coordinated transcriptional response to old age, both down- and upregulated genes within enriched functional categories show reduced expression variability and experience strong purifying selection. Collectively, our results support that the aging soma elicits a plastic transcriptional program that adjusts the organism to a declining physiological condition, implying that many age-related expression changes mitigate rather than accelerate somatic aging. These findings call for a more nuanced view of the causes and consequences of age-related transcriptional change, with implications for both theoretical and applied research on aging and disease.

## Main

Aging is thought to result from the accumulation of damage and a gradual loss of cellular homeostasis (*1*, *2*). Among the many processes that challenge homeostasis, changes in gene regulation are particularly relevant, as cellular function critically depends on finely tuned gene expression (*3*, *4*). Accordingly, the widespread expression changes observed during aging (*5–9*) have led to the hypothesis that aberrant gene expression (dysregulation) plays a key role in aging (*10–15*). While this idea certainly has some merit, it is not unreasonable to assume that many age-related expression changes instead help mitigate somatic deterioration. A complete understanding of the role of gene expression in aging therefore requires distinguishing changes that contribute to its progression from those that counteract it (*16*).

Increased dysregulation with age is a plausible scenario because gene expression is partly regulated through epigenetic mechanisms, including DNA methylation, histone modifications, and chromatin structuring (*17*, *18*). These chemical modifications are dynamic (*19–24*), and may therefore decay with age as selection to maintain their correct positioning weakens (*13*). Consistently, studies have reported that aging is linked to increased stochastic expression variability between cells (*14*, *25–28*, but see *29*), reduced coordination of expression patterns within cells (*30*), and epigenetic drift (*16*, *31*).

While dysregulation seems to increase with age, it is also plausible that many genes adjust their expression in response to declining physiological conditions associated with aging, thereby helping mitigate rather than exacerbate somatic deterioration. In accordance with this view, studies have shown age-related upregulation of repair and stress-response pathways, suggesting a coordinated regulatory response aimed at counteracting the accumulation of molecular damage (*12*, *32*, *33*). Likewise, the frequently observed downregulation of metabolic genes in older individuals (*12*, *32*, *33*) potentially serves to adjust energy expenditure to help preserve cellular and organismal homeostasis as physiological condition declines with age.

Plastic gene regulatory responses that optimize organismal function under different physiological states are generally expected to be under strong selection, because it allows individuals to maximize fitness under diverse conditions across an individual’s lifetime (e.g., with fluctuations in nutrient availability, infections, or other stressors). Condition-dependent optimization of resource use has been extensively discussed in the context of life-history evolution (*34–40*), but has largely been overlooked in studies on gene expression changes associated with aging (but see *41*). While natural selection may be too weak to fine-tune gene expression responses specifically to age-associated physiological changes, condition-dependent responses may still be induced and help slow the pace of somatic deterioration, if age-related physiological change resembles that of young individuals in poor condition. Yet whether such an overlap exists between age- and condition-related gene expression changes remains unclear.

Here, we test the hypothesis that many age-related expression changes reflect a plastic, condition-dependent response. We experimentally reduce organismal condition independently of age in female fruit flies and compare the resulting expression changes to those that naturally occur with age. Our results reveal a substantial overlap in both genes and functional categories, suggesting that many age-related changes in gene expression represent plastic responses to a soma in poor condition. Because age-related condition-dependent changes likely mitigate somatic deterioration, our findings underscore the critical need to distinguish these responses from age-related dysregulation when developing therapeutic strategies to slow or reverse aging.

## Results

To manipulate condition independently of aging, we introgressed large synthetic deletions (spanning 17-58 genes) into a population of *Drosophila melanogaster* (Figure S1) and examined their effects in heterozygous females. Deletions of this size are usually deleterious (*42*), and expected to compromise carriers’ general condition by limiting their ability to acquire and metabolize resources (*43*), thereby inducing changes in the expression of condition-sensitive genes. In total, we analysed the effects of condition (wild type vs. deletion genotypes) and age (6 vs. 35 days) across 18 different deletions (Table S1) in 286 samples of pooled female heads (Figure S1). Because each heterozygous deletion also alters the expression of genes within the deleted region and their interacting partners (*44*), we excluded these genes from our downstream analyses (see Methods and Supplementary file 1). After filtering, we identified condition-sensitive genes as those differentially expressed when all 18 deletion genotypes were analysed jointly. With this analysis, we identified three sets of differentially expressed genes (DEGs): those associated with age, condition, or both (Table S2).

Using this experimental system we (1) test for a concordant transcriptomic response between aging and experimentally reduced condition; (2) assess if these parallel responses map to a shared epigenetic gene regulatory profile; (3) use this "condition-dependent" framework to assess if loss of tissue-specific and sex-biased gene expression are caused by plastic responses rather than stochastic decay; and finally, (4) test if age-related and condition-sensitive genes show evidence of tighter, not looser, regulatory control.

### Age upregulates repair and stress responses and downregulates metabolism

We began exploring the hypothesis that aging induces a condition-dependent gene expression response by studying the age-related DEGs (young vs. old individuals). Our large sample size provided substantial statistical power and identified 8047 DEGs. Both down- and upregulated genes show gene ontology (GO) enrichment for numerous biological processes consistent with previous findings (Figure S2; Table S3). Of genes annotated with GO terms, 71% and 56% of up- and downregulated genes, respectively, show association with an enriched GO category. Downregulated genes are strongly enriched for metabolic and mitochondrial functions (e.g., oxidative phosphorylation, energy generation, amino acid and carbohydrate metabolism), cytoplasmic translation and ribosome biogenesis, as well as neuronal processes (e.g., trans-synaptic signalling and the sensory perception of smell), indicating a coordinated downregulation of energetically demanding processes, consistent with a systemic response to reduced physiological condition. The upregulated genes show GO enrichment for processes such as DNA damage response, immune activity, stress response, and wound healing. While generally attributed to loss of cellular homeostasis (*2*), these processes are also consistent with a condition-dependent stress-response.

Critically, a condition-dependent response, rather than dysregulation, would predict a clear separation of enriched GO categories between up- and downregulated genes. A previous study found that a substantial proportion of GO categories were enriched for genes both up- and downregulated with age (*31*), interpreting it as evidence of dysregulation that could disrupt mRNA stoichiometry and co-expression across the transcriptome. In contrast, our more highly replicated dataset shows minimal overlap: among 864 upregulated and 322 downregulated GO categories, only five coincide, which drops to one with a stricter P-value (0.01) threshold. These results strengthen the view that most age-related expression changes represent coordinated, condition-dependent responses rather than a global loss of regulatory control.

### Aging and reduced condition have concordant effects on gene expression

We then tested our central prediction: that age-related DEGs reflect a response to declining condition. First, we detect 493 condition-sensitive DEGs (200 upregulated and 293 downregulated, respectively), and, as predicted, these genes are highly enriched among the age-related DEGs (414 out of 493; log_2_(obs/exp) = 0.73; hypergeometric test: *P* = 3.9 × 10^-59^; Figure 1A). This enrichment is enhanced when restricting the analysis to the most significant genes (top 20% age-related DEGs: log_2_(obs/exp) = 2.37, *P* = 3.8 × 10^-82^; Figure 1B). Further refining this analysis shows stronger overlap for downregulated than for upregulated genes (Figure 1B).

**Figure 1.**
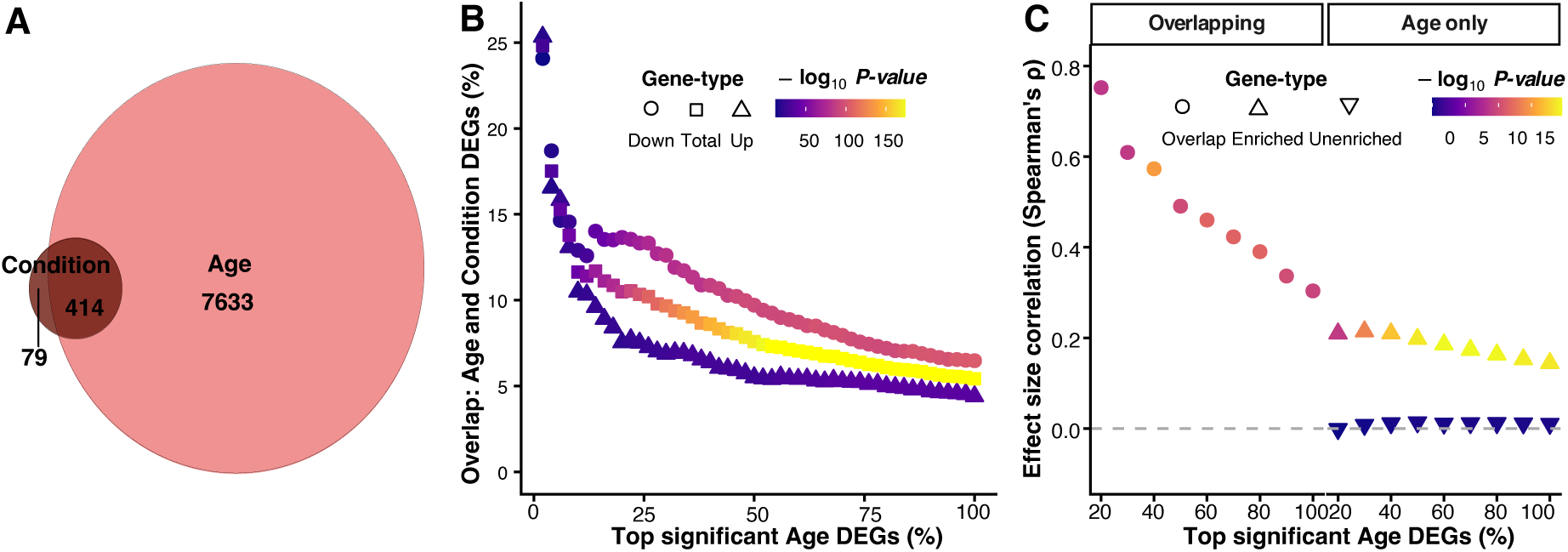
Transcriptomic responses to age and reduced condition are highly concordant. (A) Venn diagram of Age DEGs (differentially expressed genes) (35-day-old vs. 6-day-old females) and reduced Condition DEGs (females heterozygous for large deletions vs. wildtype controls). (B) Proportion of Age DEGs overlapping with Condition DEGs, as a function of the top X% most significant Age DEGs. (C) Overlapping: Spearman’s rank correlation of gene expression effect sizes (log₂ fold change) between genes DE by both Age and Condition, as a function of the top X% most significant Age DEGs. Age only: Spearman’s rank correlation for Age DEGs not DE for Condition, separated on genes within or outside Gene Ontology (GO) categories enriched for Age DEGs, as a function of the top X% least non-significant Condition genes.

Second, we find a significant positive correlation between the *effect sizes* of age and condition for the 414 overlapping genes (Spearman’s *ρ* = 0.29, *P* = 3 × 10^-10^, Figure 1C), which strengthens when the gene set is restricted to those with higher significance (top 20% most significant age-related DEGs: *ρ* = 0.75, *P* = 2 × 10^-9^, Figure 1C). With relatively few DEGs in the condition set, these correlations are nonetheless based on a small subset of genes DE with age. We reasoned that the smaller set of condition-sensitive DEGs may reflect that our condition treatment was mild compared to the effect of age (*45*) (also see limitations of this study) and may subtly affect many more genes not meeting the strict DEG threshold. We therefore tested for a correlation using genes significant only for age. Crucially, we predicted this correlation would only exist for genes in GO categories enriched for age-related DEGs (enriched genes), as only these show evidence for a coordinated response to age. Accordingly, enriched genes show a positive correlation (*ρ* = 0.14*, P* = 2 × 10^-19^) while unenriched genes do not (*ρ* = 0.01, *P* = 0.31; Figure 1C). Restricting the correlation between the effect sizes of enriched genes to those with smaller (but non-significant) P-values for condition further strengthens the correlation (top 20% genes: *ρ* = 0.21*, P* = 1.3 × 10^-9^; Figure 1C).

### Aging and reduced condition similarly modulate biological function

We next explored whether genes responding to age and condition show functional similarity by analysing the up- and downregulated condition-sensitive genes separately. We identify 58 GO categories enriched for downregulated condition genes (Table S3), showing significant overlapped with those downregulated with age compared to random expectation (25/58; Monte-Carlo permutation test: *P =* 0.03). These categories predominantly reflect metabolic processes, including organic acid, amino acid, and carboxylic acid metabolism. Within these categories, we identify several genes with potential roles in condition-dependent processes associated with mitochondrial function and a diverse array of metabolic processes. Notable examples include components of the mitochondrial pyruvate dehydrogenase (e.g., *Pdh*) and trifunctional protein complexes (*mtp*), amino acid metabolism (*bhmt, P5cr-2, P5CS, Hn, cn, Dbct, CG3999*) (*46*), lipid metabolism under dietary change (e.g., *AkhR, Mcad, Cyp6a8, walrus, lsd1*) (*47–50*) as well as feeding behaviour (*Hn, Pu*) (*48*). In contrast, only one GO category, DNA damage response, is enriched for upregulated condition genes. While this overlap is no more than expected by chance, we note that DNA damage response is the top significant GO category for upregulated age-related genes (Table S3).

To further refine our analysis, we next categorized the 200 upregulated and 293 downregulated condition-sensitive genes into six groups: four groups containing genes that show concordant and discordant changes with respect to age, and two containing genes that are up- and downregulated by condition alone, followed by separate GO enrichment analyses on each of them. DEGs showing discordant responses to reduced condition and old age, as well as those upregulated in response to condition alone, show no enrichment. Conversely, we identify 24 enriched GO categories among genes downregulated in response to both condition and age. These categories are predominantly related to metabolism and amino acid synthetic processes and show substantial overlap (17/24) with the GO categories enriched for genes downregulated by age (Figure 2A; Table S4). We also identify 7 GO categories for genes upregulated by both condition and age, of which all (7/7) are also enriched for genes upregulated by age (Figure 2A; Table S4). These categories primarily include DNA damage response and repair, and telomere/chromosome maintenance and organisation. GO categories for genes downregulated by condition alone also overlapped broadly with GO categories enriched for genes downregulated by both condition and age (8/9; Figure 2A, Table S4).

**Figure 2.**
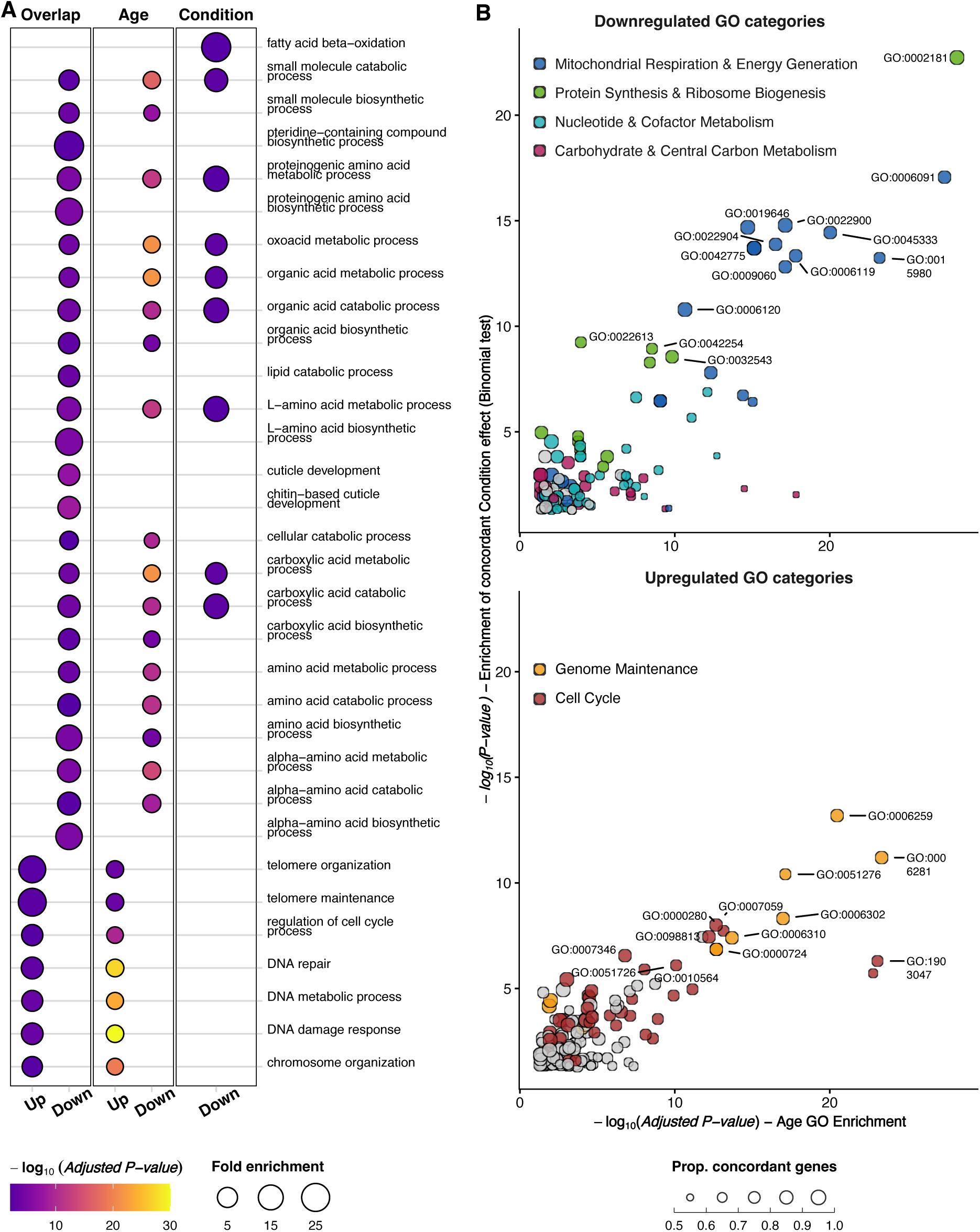
Concordance of functional enrichment between age and reduced condition. (A) Enriched GO categories for genes DE by both Age and Condition (Overlapping) and by Condition only (Condition). Additionally, the subset of GO categories that overlap with either or both of the “Overlap” and “Condition” sets when Age DEGs are analyzed separately (Age) are shown. The size of the dot represents the fold enrichment and the color bar in all plots reflects statistical significance. (B) The scatter plot displays GO categories significantly enriched for genes differentially expressed with age, and for which non-significant genes in the condition treatment show a significant concordant directional change aligning with the response to age. The x-axis represents statistical significance of GO categories enriched for significant age-related genes, and the y-axes represents the statistical significance of the same GO categories with respect to the proportion of genes showing a concordant directional change in expression in response to reduced condition. The size of the dots indicates the proportion of concordant genes within that category. Points are colored by broad biological functions. Refer to Table S5 for detailed description of GO terms.

Given the positive effect size correlation between age and condition among genes significantly affected only by age and showing GO category enrichment (Figure 1C), we hypothesized that this signal could be concentrated to specific GO categories significant exclusively for age. We therefore tested if more genes than expected by chance showed a concordant, albeit non-significant, effect of condition within each age-enriched GO category. Among the 297 categories enriched for DEGs downregulated by age (excluding the 25 that overlap with significant condition-sensitive genes), 122 show significant enrichment for a concordant condition effect (Table S5), far exceeding random expectation (mean = 10.4; median = 8; *P* < 7 × 10^-5^). In addition to providing further evidence for downregulation of metabolism, this analysis provides strong support that reduced condition downregulates genes in top age-associated GO categories for downregulated genes, including mitochondrial function, cytoplasmic translation, and ribosomal gene expression (Figure 2B; Table S3; Table S5). Notably, the downregulation of ribosomal gene expression under reduced condition aligns with a previous study that identified it as the most conserved signal associated with aging across cell types in *Drosophila* (*9*). Similarly, of the 863 GO categories enriched for upregulated age-related genes, 160 show significant enrichment for genes with concordant condition-effect (Table S5), also exceeding random expectation (mean = 27.9; median = 23; *P* = 2 × 10^-4^). This analysis provides additional support for upregulated genome maintenance (DNA repair and organisation, as well as chromatin and telomere maintenance and organisation), but also for upregulated of cell cycle processes (Figure 2B). This latter result may seem counterintuitive at first, as head-tissue cells are post mitotic. However, recent data show that polyploid cells accumulate in terminally differentiated neurons and glia, and that polyploidy protects against DNA damage-induced cell death (*51*). This result thus provides further support for upregulation of damage control when condition is reduced.

### Age-related and condition-sensitive genes share a histone methylation profile

We hypothesized that these plastic, enriched genes would be regulated by mechanisms sensitive to physiological state, such as the tri-methylation on histone H3 lysine 4 (H3K4me3), an activating histone mark (*52*), which has been shown to respond to nutritional status (*53*, *54*). Specifically, genes marked with H3K4me3 in young flies could be downregulated with age through reduced methylation intensity. While such changes could occur through stochasticity, a plastic response would result in H3K4me3 associations being more common among enriched genes than among unenriched genes or genes that do not change expression with age (i.e., the ‘non-significant’ genes). We tested this possibility using female head tissue H3K4me3 data from Nanni et al (2023) (*55*). Consistent with this hypothesis, we find that downregulated age-related enriched genes are more frequently associated with H3K4me3 than their unenriched counterparts (86.7% vs. 71.5%, *χ*^2^ = 118.72, *df* = 1, *P* = 1.2 × 10^-27^; Figure 3A), as well as the non-significant genes (44.6%; enriched: *χ*^2^ = 879.02, *df* = 1, *P* = 3.6 × 10^-193^; Figure 3A). Importantly, genes downregulated by condition are also more frequently associated with H3K4me3 than genes non-significant with respect to condition (67.6% vs. 61.5%, *χ*^2^ = 4.09, *df* = 1, *P* = 0.04; Figure 3A). Because our condition treatment may affect not only condition-sensitive genes, but also genes with relatively unstable expression patterns, we also compared downregulated condition genes enriched for a GO category with genes whose expression was unchanged by the treatment. The point estimate for the association with H3K4me3 is higher for enriched downregulated condition genes (71.6%), but not significantly different from the non-significant genes (*χ*^2^ = 3.61, *df* = 1, *P* = 0.06).

**Figure 3.**
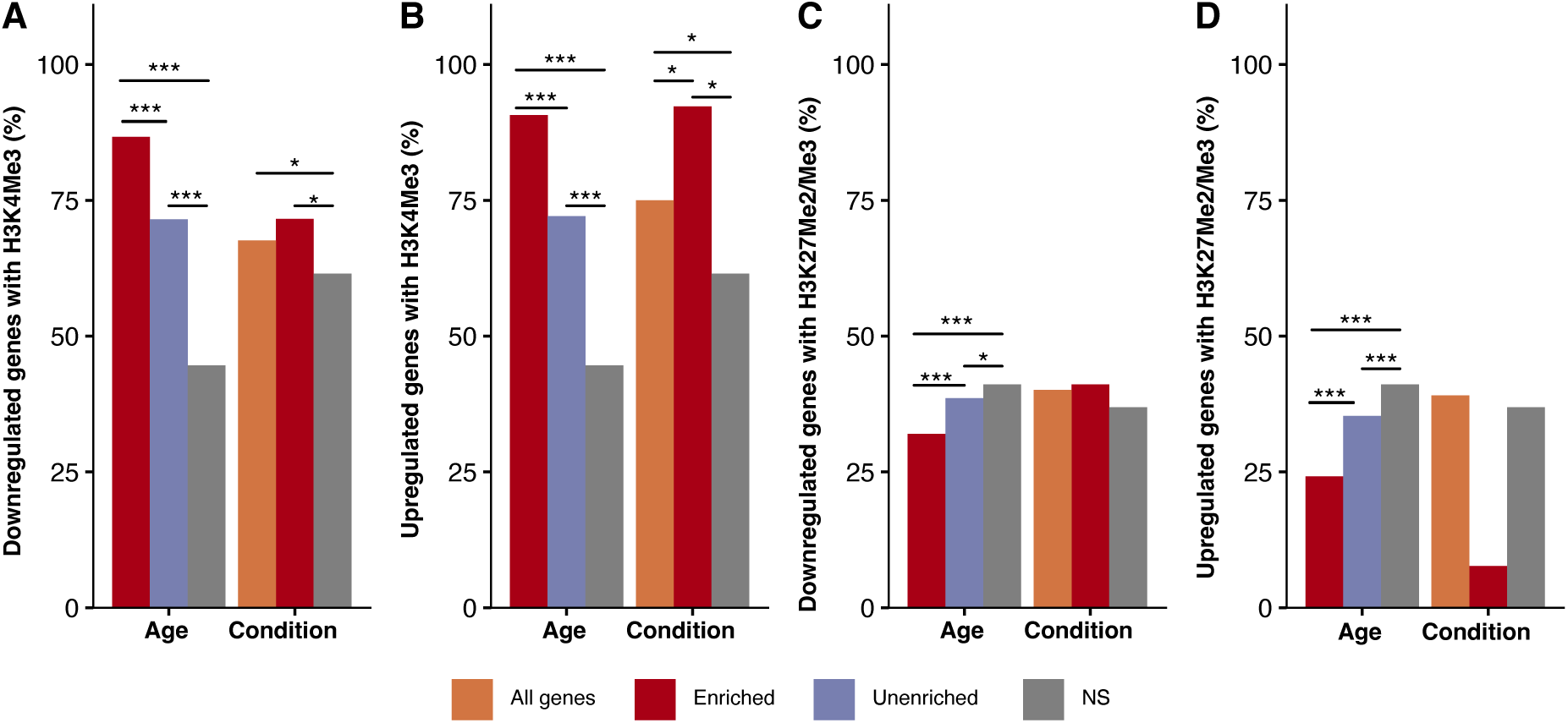
Enriched DEGs are associated with epigenetic marks of active gene regulation. Bar plots show the percentage of downregulated (A, C) and upregulated (B, D) genes associated with the activating histone mark H3K4me3 (A, B) or the repressive mark H3K27me2/me3 (C, D). Genes are categorized as DEGs for Age or Condition. Age DEGs are further subdivided into those belonging to a functionally enriched Gene Ontology category (Enriched; red) and those that do not (Unenriched; blue). All condition genes (all genes; orange) were further refined into enriched condition genes. The frequency in genes not significantly affected by the respective treatment is shown as a baseline (NS, grey). Statistical significance is denoted by *** (P<0.001) and * (P<0.05).

Previous studies have indicated that genes involved in the DNA damage response are upregulated via increased H3K4me3 levels (*56–58*), and we reasoned that other genes with stress-related functions might also be upregulated through this mechanism. Supporting this idea, we find that upregulated age-related enriched genes show a stronger association with H3K4me3 than unenriched genes (90.7% vs. 72.1%, *χ*^2^ = 229.4, *P* = 8.0 ×10^-52^; Figure 3B), and that this pattern is further stronger when the comparison in made to the non-significant genes (44.6%; enriched: *χ*^2^ = 1420.4, *df* = 1, *P* = 7.5 × 10^-311^; Figure 3B). Genes upregulated by reduced condition also show higher association with H3K4me3 than genes that do not change expression with condition (75% vs. 61.5%, *χ*^2^ = 14.0, *df* = 1, *P* = 1.8 × 10^-4^; Figure 3B), and this difference is further strengthen when only enriched upregulated condition genes are considered (92.3%; *χ*^2^ = 3.97, *df* = 1, *P* = 0.046). Together, these results show that both up- and downregulated enriched age-related genes have stronger associations with H3K4me3, a pattern that is also seen in condition-sensitive genes. This suggests that changes in H3K4me3 marking help regulate expression shifts in enriched age-related genes as a plastic response to age-related changes in condition.

In contrast to H3K4, methylation of H3K27 is associated with transcriptional repression(*59*). The regulatory role of H3K27 methylation is less well understood, but it has been suggested that age-related epigenetic drift of H3K27me3 may contribute to gene dysregulation associated with aging (*60*). Using head tissue H3K27me2me3 data from Nanni et al. (2023) (*55*), we find that among genes both down- and upregulated regulated by age, enriched genes show reduced association with H3K27me2me3 compared to unenriched (downregulated: 32.0% vs 38.6%, *χ*^2^ = 16.0, *P* = 3.8 × 10^-5^; Figure 3C; upregulated: 24.2% vs 35.3%, *χ*^2^ = 57.6, *P* = 3.1 × 10^-14^; Figure 3D), as well as compared to non-significant genes (41.1%, downregulated: *χ*^2^ = 43.9, *P* = 3.4 × 10^-11^; Figure 3C; upregulated: *χ*^2^ = 204.4, *P* = 2.3 × 10^-46^; Figure 3D). Genes down-(40.1%) or upregulated (39.1%) by condition are, however, not differently associated with H3K27me2me3 compared to non-significant genes (36.9%; P > 0.05; Figure 3D). Focusing only on enriched genes, this conclusion holds for both down- and upregulated genes (41.1% and 7.7%, respectively; both *P* > 0.05). Together, these results specifically link the plastic changes in both age and condition to the dynamic H3K4me3 mark, while the repressive H3K27 mark appears disassociated exclusively with enriched age-related genes.

### Loss of tissue identity with age is a condition-dependent response

“Loss of tissue identity”, the downregulation of tissue-specific genes with age, is often interpreted as regulatory decay (*14*, *61–66*). However, a plastic, condition-dependent response to conserve energy by deprioritizing tissue-specific tasks would generate the same pattern. To test this hypothesis, we first calculated a head-biased expression value (*z*-score) for each DEG, using expression data across female tissues from FlyAtlas2 (*67*). We then classified DEGs with high positive or negative *z*-scores as either head-biased (> 1.5) or reverse-head-biased (< -0.5). Using this data, we confirm that age-downregulated genes are enriched for head-biased genes compared to upregulated ones (30.6% vs. 15.1 %; *χ*^2^ = 254, *P* = 2.56 × 10^-57^; Figure 4A). Consistent with our hypothesis, genes downregulated by condition show the same pattern (38.7% vs 28.1 %; *χ*^2^ = 5.0, *P* = 2.5 × 10^-2^; Figure 4A). Age-related genes with reverse-head-biased expression show no difference between down- and upregulation (3% for both, *P* = 0.47; Figure 4A), which also is the case for condition genes (1% vs. 3 %; *χ*^2^ = 3.7, *P* = 0.054; Figure 4A).

**Figure 4.**
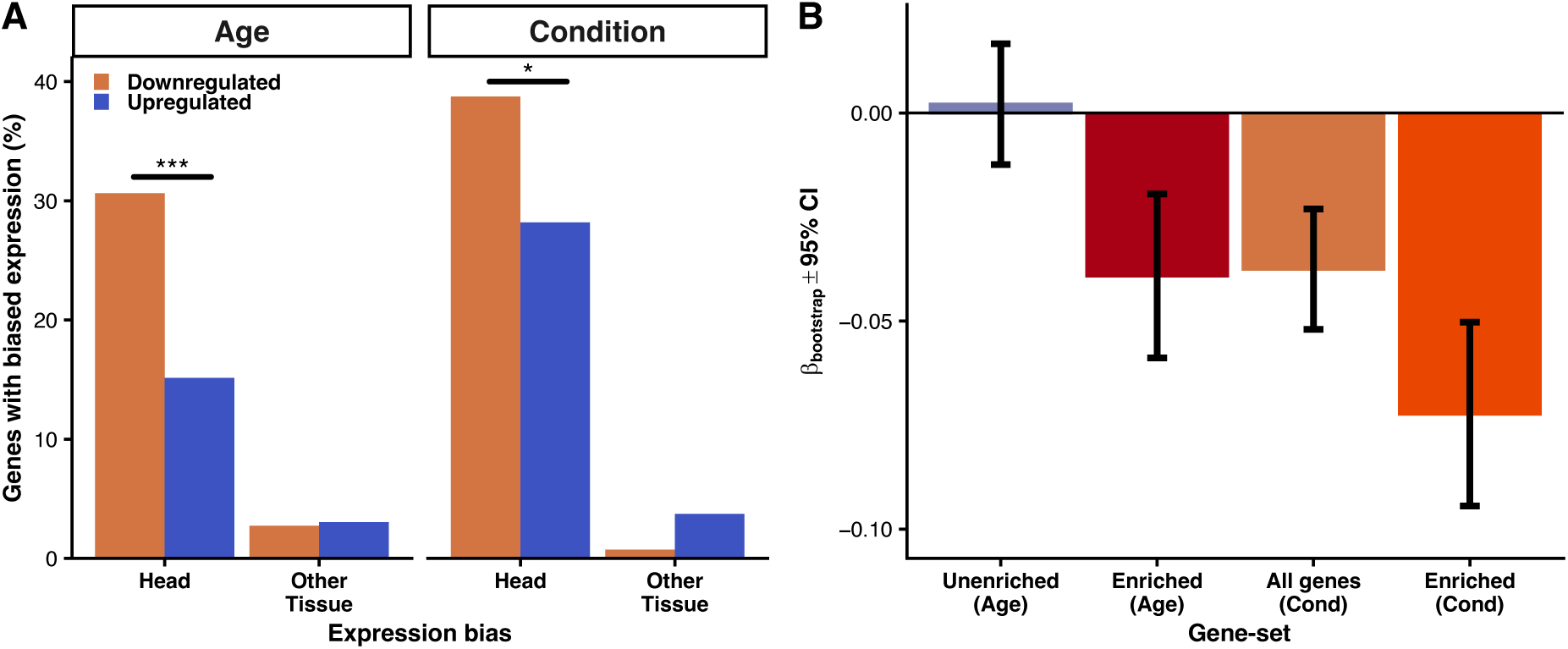
Age-related loss of tissue identity is caused by condition-dependent downregulation. (A) Percentage of upregulated and downregulated DEGs for Age and Condition that are classified as head-biased (Head, i.e., having preferentially high expression in the female head tissue) or other tissue biased (Other Tissue, i.e., having preferentially high expression in a female tissue other than the head). Significance is denoted by *** (P<0.001) and * (P<0.05). (B) Bootstrap regression analysis showing the relationship between the degree of head-biased expression (z-score) and the magnitude of gene downregulation (log₂-fold change). Bootstrap regression coefficients (βbootstrap) and associated 95% confidence intervals are displayed.

To further investigate the cause to loss of tissue identity with age, we tested for a negative association between the degree of head-biased expression and age-related gene downregulation. Both the condition and the dysregulation hypothesis predict a negative association, but we reasoned that a stronger negative association among enriched than among unenriched genes would support the condition hypothesis. Using a bootstrap approach, we find that, among genes downregulated by age, the bootstrap regression coefficient (*β*_bootstrap_) is significantly negative for the enriched genes (*P =* 5 × 10^-3^; Figure 4B), but not for the unenriched ones (*P* = 0.57), supporting the condition hypothesis. This conclusion gains further support when we analyse downregulated condition genes, as also these show a negative association between the degree of head-biased expression and downregulation (*P =* 0.03; Figure 4B). The slope is further significantly steeper when we restrict the analysis to enriched condition-sensitive genes, as compared to all condition-downregulated genes (Δ*β*_bootstrap_ = 0.04, P <0.0001; Figure 4B).

### Loss of sex-biased expression with age is a condition-dependent response

Sex-biased expression can be viewed as a special form of tissue-specific expression and, consistent with the condition hypothesis, it has previously been shown to decline under reduced condition (*68*, *69*). To test whether this pattern also emerges during aging, and in response to our condition treatment, we examined expression changes in a set of 540 female-biased (FB) genes identified in brain tissue from young females, from a study based on the same population as used here (*41*). Supporting the condition hypothesis, these FB genes are significantly overrepresented among downregulated rather than upregulated genes DE by age (log_2_[obs/exp] = 1.4, *χ*^2^ = 97.4, *P* = 5.4 × 10^-23^; Figure 5), and this pattern is even stronger for condition DEGs (log_2_[obs/exp] = 3.3, *χ*^2^ = 41.4, *P* = 1.24 × 10^-10^; Figure 5). Furthermore, this signal is strengthened when the comparison is restricted to enriched downregulated genes for both age (hypergeometric test: *P =* 0.04) and condition (hypergeometric test: *P* = 0.03; Figure 5). Finally, FB genes are significantly overrepresented among genes downregulated by both age and condition (*P* = 4.6 × 10^-5^). Together, these findings suggest that, like tissue-biased genes, the age-related decline in female-biased expression reflects a coordinated, condition-dependent response.

**Figure 5.**
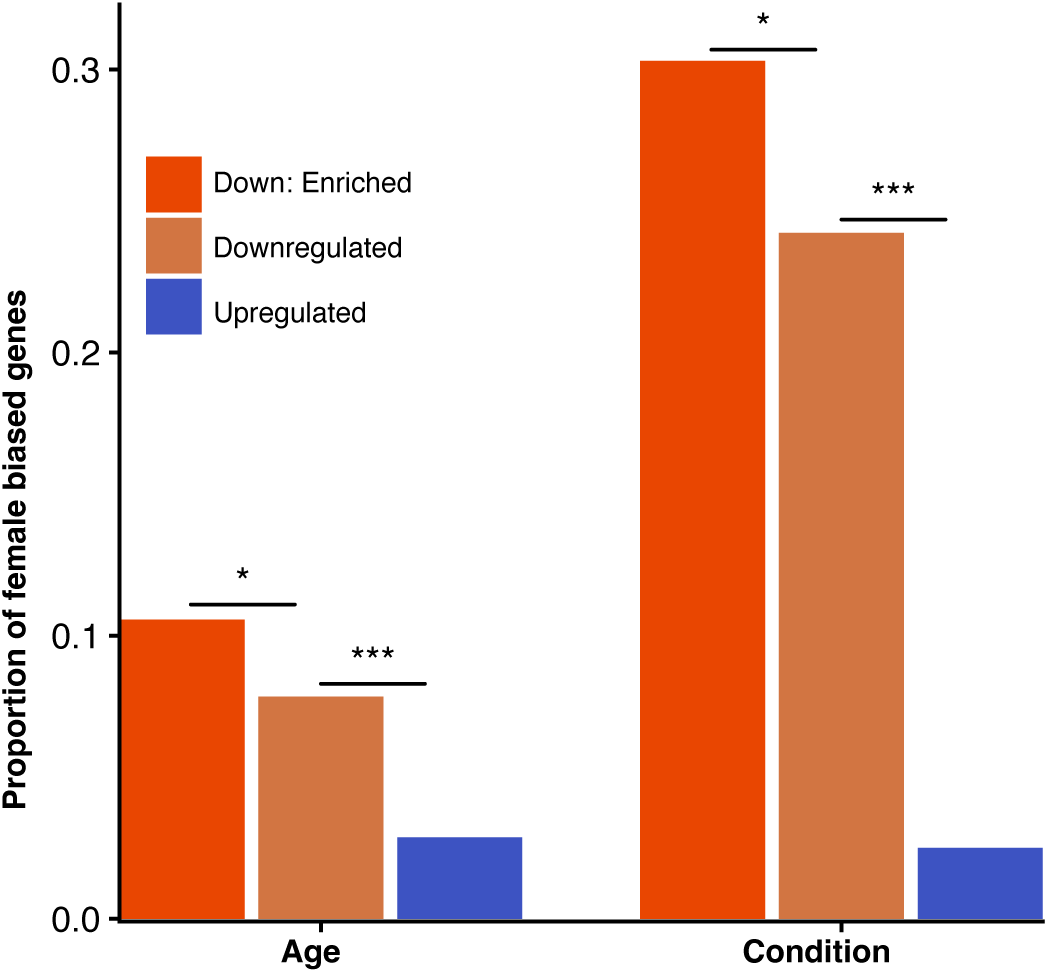
Female-biased gene expression declines with age and reduced condition. The proportion of female-biased genes is shown for different sets of differentially expressed genes. Genes are categorized as upregulated (blue) or downregulated (orange) by Age or Condition. The downregulated sets are further refined into genes belonging to an enriched GO category (Down: Enriched; red). Statistical significance denoted by *(P<0.05), *** (P<0.001).

### Strong regulatory constraint on functionally enriched genes

If many genes change expression with age as a condition-dependent response rather than regulatory breakdown, we expect them to be tightly regulated. This possibility would be supported if genes in enriched GO categories experience stronger stabilizing selection on their expression than unenriched genes. While we cannot directly test this prediction, we reasoned that the strength of stabilizing selection acting on a gene should be reflected by its expression variability across samples. To assess this, we calculated the coefficient of variation (CV) of the DEGs in both young and old samples and compared enriched to unenriched genes. Consistent with our prediction, CVs are lower for enriched downregulated genes in both young (*P* = 6 × 10^-28^; Figure 6A; Table S6) and old (*P* = 8 × 10^-31^; Figure 6A; Table S5) samples, a pattern that also holds for upregulated genes (*P* = 8 × 10^-36^ and 9 × 10^-50^ for young and old, respectively; Figure 6A; Table S6).

**Figure 6.**
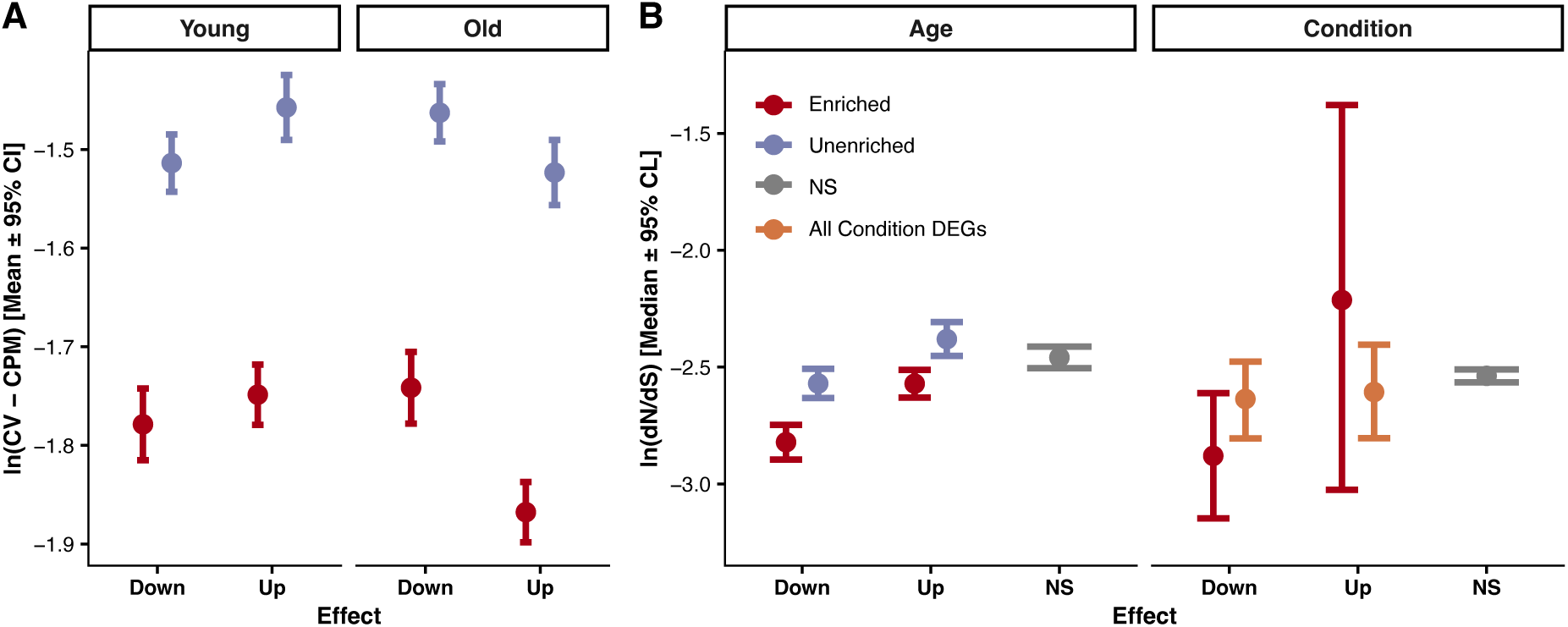
Enriched DEGs exhibit signatures of tight regulatory control and strong purifying selection. (A) Expression variability, measured as the log-transformed coefficient of variation (CV) of counts-per-million (CPM), is shown for different gene sets in young and old samples. (B) Evolutionary constraint, measured as the ratio of nonsynonymous to synonymous substitution rates (dN/dS). A lower dN/dS value indicates stronger purifying selection. Genes categorized as Enriched (red); Unenriched (light blue); Condition (orange); Non-Significant (NS) genes (grey). In both panels, error bars represent 95% confidence/credibility intervals.

Previous studies have found that stabilizing selection on gene expression is often positively correlated with the strength of purifying selection on coding regions(e.g., *70*–*76*) (but see (*77*)). We therefore reasoned that evidence of stronger purifying selection acting on enriched genes would further support the scenario where these genes are under stronger stabilizing selection. To assess this, we first examined the tentative link between stabilizing and purifying selection by testing for a correlation between the CV and dN/dS (the ratio of nonsynonymous to synonymous divergence). Across all genes, the correlation is significantly positive, whether the CV was estimated from young (Spearman’s *ρ* = 0.18, *P =* 2.7 × 10^-63^) or old (*ρ* = 0.15, *P =* 2.1 × 10^-43^) flies. We next compared dN/dS values between enriched and unenriched genes using Bayesian regression models. Enriched genes have significantly lower dN/dS values than their unenriched counterparts for both down- and upregulated genes (Table S7 Figure 6B).

Enriched down- and upregulated genes, as well as unenriched downregulated genes, further have lower dN/dS values than non-significant genes, while unenriched upregulated genes show no difference from this group (Table S7, Figure 6B). To conduct a more detailed comparison to the non-significant genes, we first conducted a separate GO enrichment analysis on these genes and then separated them into an enriched and unenriched group. Significantly downregulated enriched and unenriched genes both have lower dN/dS values than their non-significant counterparts, while the same comparisons between significant upregulate genes and non-significant genes show no difference (Table S7). We also observe lower, albeit statistically non-significant point estimates of dN/dS for both down- and upregulated condition-sensitive genes than for non-significant genes (Table S7, Figure 6B). Restricting the above analysis to enriched genes shows significantly lower dN/dS values for downregulated, but not for upregulated genes (Table S7, Figure 6B).

## Discussion

The widespread expression changes associated with age are not uncommonly interpreted as dysregulation contributing to aging. While this view may be partly correct, we present several lines of evidence indicating that many of these changes instead reflect plastic responses to a soma in declining physiological condition, potentially helping to mitigate rather than exacerbate somatic deterioration. First, among genes responding to the experimentally reduced condition, the vast majority overlap with those that change expression with age, and age- and condition-related expression changes are positively correlated. Second, GO categories enriched for genes responding to reduced condition largely overlap with those enriched for genes responding to age, and these categories primarily reflect an expected downregulation of metabolism, energy production and protein synthesis, alongside upregulation of genome maintenance. Third, age-related genes in enriched GO categories and condition-sensitive genes appear to share a regulatory mechanism. Fourth, tissue- and sex-biased genes are downregulated by both condition and age, consistent with the idea that reduced condition leads to a deprioritizing of specialized functions in favour of core cellular processes. Finally, we find that enriched age-related and condition-sensitive genes are under strong selective constraints, suggesting that they remain tightly regulated and change expression with age as a coordinated response to reduced condition rather than from dysregulation.

### Condition and age influence overlapping sets of genes and functional categories

GO categories enriched for genes upregulated with age in this study overlap with those consistently identified across tissues (*33*, *78*) and taxa (*12*, *16*, *32*). These primarily include genes involved in various stress responses, such as immune and inflammatory processes, and in genome maintenance, which is unsurprising given the diverse types of damage expected to accumulate with age (*1*, *79*). For genes upregulated by reduced condition, GO enrichment represents a subset of the age-related categories and is confined to DNA damage response and organisation. The concerted directional bias among non-significant condition-sensitive genes within genome maintenance GO categories further reinforces increased investment in genome maintenance under reduced condition. Whether reduced condition causes genotoxic stress or a proactive resource shift to somatic maintenance is unclear. For example, a mild reduction may not be enough to trigger a stress response but could still upregulate stress-response genes by shifting allocation of resources towards somatic maintenance. Supporting this, dietary restriction in mice increases the DNA damage response (*80*, *81*), although this has not been reported for *Drosophila* (*82*). Conversely, a more severe reduction may impair core cellular functions and thereby trigger various stress responses. In any case, we conclude that the stress response to our condition treatment was mild compared to that induced by age.

Genes downregulated with age across taxa and tissues are primarily associated with metabolism and mitochondrial function, along with protein synthesis and turnover, growth signalling, and developmental processes (*12*, *16*, *32*, *33*, *78*). We similarly detect a clear downregulation of metabolism, mitochondrial function, and protein synthesis and turnover with age. However, growth- and development-related categories are unexpectedly enriched for upregulated genes. GO categories enriched for genes that are downregulated in response to reduced condition are predominantly associated with metabolism and largely overlap with those affected by age. The overlap is primarily focused to amino acid, organic acid, carboxylic acid, and small-molecule metabolism, while lipid metabolism is more pronounced in the condition treatment. These results are consistent with the expected shift away from resource-intensive processes when condition is reduced, a pattern also observed in *D. melanogaster* when adult nutrient access is restricted (*83*). Several genes involved in mitochondrial function and protein turnover are also significantly downregulated under reduced condition. However, downregulation of genes with these functions under reduced condition becomes strikingly apparent when the direction of change of non-significant genes is considered, as this closely parallels the age response of these genes. This coordinated suppression across genes involved in mitochondrial function and protein turnover indicates that reduced condition induces a subtle yet systematic transcriptomic shift that recapitulates key features of aging. Intriguingly, condition-induced decline in mitochondrial activity may nuance the concept of mitochondrial dysfunction as a hallmark of aging (*2*, *84*), by suggesting that at least some of the age-related decline in mitochondrial function reflects a regulated response to a somatic state of diminished physiological condition, rather than degeneration.

### Condition and age downregulate tissue- and sex-biased genes

While many genes alter their expression with age, relatively few change concordantly across tissues (*9*, *33*, *85*). At the tissue level, this pattern can be partly explained by age-related loss (or gain) of tissue-specific cell types (*65*), but downregulation of genes with tissue-specific expression also contributes, often interpreted as stochastic loss of tissue-specific regulatory mechanisms (*61*, *63*, *64*). Here, we also observe downregulation of tissue-specific genes with age. Because *Drosophila* head tissue shows little age-related loss of tissue-specific cell types (*9*), this likely reflects a genuine transcriptional decline. Crucially, we find a similar downregulation of tissue-specific genes under reduced condition. This finding does not support the idea that downregulation of tissue-specific genes with age reflects regulatory breakdown. A more parsimonious interpretation is a plastic response to a soma in poor condition, where tissue-specific tasks are deprioritized in favour of core cellular processes. This conclusion gains further support from our finding that female-biased genes are downregulated both with age and reduced condition, as investments in the expression of sex-biased genes, as well as sexual traits, relate to general condition (*36*, *68*, *69*, *86–88*).

### Condition- and age-sensitive genes share an epigenetic regulatory mechanism

Additional support for the idea that many age-related expression changes reflect a response to a soma in poor condition comes from our finding that age-sensitive genes in enriched GO categories and condition-sensitive genes share enrichment for H3K4me3, a histone mark associated with active transcription (*52*). This result suggests that condition-dependent signalling may modulate their expression through altered H3K4me3 marking. However, shared H3K4me3 enrichment does not necessarily imply coregulation, as stochastic gain or loss of H3K4 methylation can generate similar changes in expression patterns. Two observations argue against this interpretation: first, the turnover of H3K4me3 is rapid (*89*, *90*), making persistent stochastic effects unlikely; second, dynamic age-related H3K4me3 modifications primarily occur at marks acquired during adulthood rather than development (*91*), consistent with context-dependent regulation rather than random chromatin deterioration.

Studies in nematodes have identified a connection between lifespan and genes regulated by H3K4 methylation, in which reduced expression of H3K4me3 methyltransferases, such as set-2, extends lifespan (*92–94*). These methyltransferases utilize S-adenosylmethionine (SAM) as a methyl donor, and SAM availability is influenced by the methionine cycle, which is tightly coupled to nutritional status (*53*, *54*, *95*). Lower nutritional status, resulting from aging or reduced condition, could thus lead to the observed downregulation of metabolic genes through restricted SAM-dependent methylation, and cause reallocation of limited resources towards somatic maintenance. This mechanism, however, cannot directly explain the concurrent enrichment of H3K4me3 among genes upregulated by age and reduced condition. These genes are primarily involved in various stress responses, and previous studies suggest that H3K4me3 marking of such genes can facilitate their activation under stressful conditions by broadening or intensifying H3K4me3 domains (*96*, *97*). The molecular basis of this regulatory process remains unclear.

### Strong selection on age-sensitive genes

As a final piece of evidence supporting the hypothesis that many genes change expression with age as a coordinated response rather than becoming dysregulated, we find that downregulated age-related genes in enriched GO categories are subject to stronger purifying, while upregulated age-related enriched genes are under no weaker purifying selection than other genes. A more stringent test of the hypothesis that genes change expression with age as a plastic response would demonstrate that the expression of these genes is under stronger stabilizing selection. Our finding that these genes exhibit reduced expression variation between samples is indeed consistent with this prediction, which follows if they have reduced genetic variation in expression. However, an increased robustness of these genes to minor environmental perturbations could also explain reduced variability among samples.

### Implications for theories of aging

Our results on the strength of purifying selection across gene classes challenge how similar findings have previously been interpreted in the context of evolutionary theories of aging. Several earlier studies have found that genes upregulated with age are under less evolutionary constraint than downregulated ones (e.g., *98–101*, but see *33*). This pattern has been viewed as supporting the Mutation Accumulation theory of aging, which proposes that aging evolves because the declining strength of selection with age causes increased accumulation of deleterious mutations in late-acting genes (*102–104*). While our results confirm that upregulated genes are, on average, under less evolutionary constraint than downregulated genes, they add a critical nuance: upregulated genes are not under weaker purifying selection than non-age-related genes and, display the lowest expression variation compared to all other gene categories in old samples. Moreover, most upregulated genes (71%) fall into enriched GO categories, many of which are associated with genome maintenance and stress response processes. These findings do not support the idea that upregulated genes are replete with deleterious mutations that cause aging but instead favour the view that many of these genes help mitigate somatic decline. This conclusion is further reinforced by the observation that upregulated genes can face even stronger purifying selection than genes downregulated with age, as is the case in highly proliferating mammalian cancer-prone tissues (*101*).

Although the condition-dependent up- and downregulation of gene expression we observe at old age likely slows somatic decline, these changes are not necessarily adaptive from a fitness perspective. As aging advances, there may come a point at which terminal investment into reproduction yields a higher fitness payoff than continued somatic maintenance (*105*). Investments in maintaining the soma beyond this point could therefore actually accelerate aging, if organismal aging is view from holistic perspective taking also reproduction into account. However, evolving an optimal switch point between maintenance and terminal investment is challenging, since selection may be unable to distinguish reliably between a temporary and an irreversible decline in condition. Moreover, selection acting at later ages may be too weak to drive such a switch. The condition-dependent plastic response we observe may therefore represent Antagonistic Pleiotropy (*106*): the same response that presumably is adaptive early in life, when recovery from reduced condition is possible, may become maladaptive later, when aging has reduced condition beyond the point of recovery.

### Limitations from manipulating condition through deletions

We manipulated condition using heterozygous deletions that directly alter the expression of numerous genes, disrupting regulatory networks and presumably reducing physiological condition through diverse cellular processes. We defined condition-sensitive genes as those affected when all 18 deletions are modelled jointly. This approach, however, likely underestimates the true overlap between aging and condition-dependent responses for several reasons. First, the deletions probably overlap only partially in the cellular processes they affect, resulting in heterogeneity across deletions and less comprehensive perturbations than those caused by aging. Consistent with this, genetic and environmental manipulations of condition do not perfectly align (*38*, *107*). Many condition-sensitive genes may therefore have responded to only some deletions, producing a restricted set of significantly affected genes, a possibility supported by the positive correlation across treatments for genes exclusively significant for age. Second, the milder impact of deletions compared to aging, as indicated by their much smaller effect on fecundity (*45*), likely further reduced statistical power. Third, because nonlinear expression changes with age are common (*108*), the different magnitudes of condition reduction through deletions and age can further have complicated interpretation. Indeed, genes downregulated exclusively by the condition manipulation and genes downregulated by both age and condition share most GO categories (Figure 2A). Fourth, some easily perturbed genes may have responded to the condition manipulation without being condition-sensitive, diluting the signal from true condition-sensitive genes, a possibility supported by the stronger results often observed when analyses were restricted to condition genes showing GO enrichment. Finally, a general mechanism buffering the expression of genes with altered copy number could exist. Such a mechanism would cause genes to respond to all deletions without being truly condition-sensitive and weakening results. Evidence for such a buffering system is, however, limited (*44*) and the deleted genes studied here were on average only marginally buffered (data not shown).

### Conclusions

The power of epigenetic clocks to predict biological age (e.g., *109*) has given the impression that accumulating stochastic changes to the epigenome gradually erode regulatory control and drive many age-associated expression changes. However, the recent finding that clock values can tick backwards when individuals recover from temporary condition-reducing events such as surgery, pregnancy, or serious infection (*110*) calls this interpretation into question. It suggests that many age-related epigenetic and transcriptional changes are instead plastic responses to declining physiological condition. Consistent with this view, a large proportion of the age-associated expression changes we observe are functionally enriched and overlap with responses to poor condition, suggesting that many expression changes related to genome maintenance, metabolism, mitochondrial function, and protein turnover likely mitigate rather than exacerbate somatic deterioration. This insight has implications for interventions aimed at rejuvenating organisms by reverting epigenetic marks and gene expression to youthful states. Correcting genuinely dysregulated genes should indeed help restore cellular functions. In contrast, reversing adjustments that mitigate damage or optimize resource allocation for the current somatic state could be harmful if the underlying causes of poor condition are not simultaneously addressed.

## Materials and Methods

### Fly population

We used flies from a laboratory-adapted population of *Drosophila melanogaster* known as Dahomey. This population was collected from the wild in Benin (previously Dahomey) in 1970 (*111*). It has, over the last 50 years, been kept in cages as a large, outbred population with overlapping generations, with bottles of food placed in the cages every week and cycled out after 4 weeks. The population has been kept under constant conditions (standard yeast/sugar-based food medium, 12-h light:12-h dark cycle, 25 °C, and 60% relative humidity). The overlapping generations, the large population size (not controlled but in the thousands) and the controlled conditions makes the Dahomey population a well-suited model for studies on longevity and aging, and it has become widely used in this research field, e.g., (*112–121*).

### Experimental Procedures

We backcrossed 18 autosomal genomic deletions from the Exelixis deletion collection (*122*), spanning 17-58 genes, into a copy of the Dahomey population (Dahomey-*w*, which we had made homozygous for *white* (*w*), a recessive eye colour marker), for at least eleven generations (see Figure S1 and Brengdahl et al 2023 for details). We first searched FlyBase (May 14, 2018) for genes with effects on “Ageing”, “Aging”, “Senescence”, “Lifespan”, and “Life span”, and excluded any deletion spanning genes associated with one or several of the search terms. To avoid testing the same gene multiple times, we included only deletions spanning mutually exclusive segments of the genome. Deletions were obtained from the Bloomington Drosophila Stock Center.

To reduce environmental variation affecting the experimental flies, we cultured each deletion line at a controlled density (∼180 eggs per vial) for two generations prior to the experiment. In the second generation, males heterozygous for the deletion were crossed to Dahomey-*w* virgin females and daughters heterozygous for the deletion and wild-type daughters from this cross were used in the experiment. This setup, in which mothers of all focal wild-type and deletion females share the same genotype, avoids maternal effects influencing results both within and across deletion lines. Since the same males fathered both mutant and wild-type females within a deletion line, our design also avoids potential paternal effects (*123*, *124*).

Ten days after egg laying, we collected the experimental flies and housed 33 deletion females with 33 wild-type females and 33 Dahomey*-w-e* males per experimental vial (Figure S1). The recessive body colour marker *ebony* (*e*) had been introgressed into the Dahomey-*w* background at an earlier stage, and males from this stock were used to enable quick sorting of females from males under CO_2_ anaesthesia. Throughout the experiment, flies were transferred to vials with fresh food four times per week. Dahomey*-w-e* males were replaced at days 13 and 27 with ∼4-day-old males to standardize the level of harassment/courtship/mating the females experienced as they aged. To further standardize the environment for wild-type and deletion females, all flies were kept in constant darkness throughout the experiment (except during handling). This precaution was taken since the deletions are flanked with *w^+mC^*, a gene that partly restores eye pigmentation of deletion flies (to make it possible to distinguish them from wild-type flies) and thus provides deletion flies with better visual acuity. To obtain samples of young flies, five-day-old wild-type and deletion females from 4 separate vials were separated from each other (and males) under CO_2_ anaesthesia and stored in individual vials. The following day, these flies were flash frozen in liquid nitrogen. To obtain samples from old flies, 33-day-old females were separated two days before flash-freezing.

### Tissue dissection and RNA isolation

Samples of young and old flies were flash frozen at the same time of day, and any dead flies were removed before freezing. Frozen samples were stored at -80°C. Heads were subsequently dissected under a stereomicroscope in RNAlater (Thermo Fisher) and pooled in groups of 20 to reduce environmental variation (*125*). Four independent biological replicates per age, genotype, and deletion line were collected (2×2×18×4 = 288 samples in total), and stored in RNAlater at -80°C until RNA extraction.

Total RNA was extracted using an Agencourt RNAdvance Tissue kit (Beckman Coulter Inc., USA) in the tube format, following the manufacturer’s standard protocol. The samples were quantified using a Nanodrop 2000c spectrophotometer (Thermo Scientific, USA) and then quality checked with a Bioanalyzer 2100 (Agilent, USA) using an RNA 6000 Nano kit. All samples had RNA concentration above 21 ng/µL. Final RNA samples were resuspended in nuclease-free water and stored at −80 °C until library preparation.

### Library preparation and RNA-Seq

Libraries were prepared using the Illumina TruSeq Stranded mRNA Library Prep Kit (Illumina Inc., USA) according to the manufacturer’s protocol. Paired-end sequencing of mature RNA was performed at National Genomics Infrastructure Sweden on NovaSeq6000 (NovaSeq Control Software 1.7.5/RTA v3.4.4; Novogene) sequencer using ’NovaSeqXp’ workflow in ’S4’ mode flow cell. FastQ files for quality check were generated using bcl2fastq_v2.20.0.422 from the CASAVA software suite. The quality scale used was Sanger/phred33/Illumina 1.8+.

Quality control and processing of raw reads to gene counts were done using the NF-core/rnaseq workflow version 3.0 (https://nf-co.re/rnaseq) (*126*). Briefly, initial quality control was performed using FastQC v 0.11.9 (*127*). Raw reads were quality-filtered, and adaptors were trimmed using Trim Galore 0.6.6 (*128*). Reads were mapped to the *D. melanogaster* reference genome (BDGP6) using the Star aligner version 2.6.1d(*129*). Gene and transcript quantification was performed using Salmon v1.4.0 (*130*). Further details about the steps in the NF-core/rnaseq workflow are available on GitHub (https://github.com/nf-core/rnaseq). All samples had >80% of reads mapping to unique genomic positions, and only samples with more than 20 million read count were retained for downstream analyses, resulting in 286 total samples (72, 71, 71,72 for young wildtype, young mutant, old wildtype, old mutant, respectively).

### Analysis of gene expression data

After applying several filtration criteria (see Supp file S1 for details), 15851 genes were retained for analysis. Because many genes are thought to undergo modest changes in expression with age, a large proportion of differentially expressed genes are believed to go undetected in studies with smaller sample sizes (*41*, *131*, *132*). Our dataset with a large sample size (N = 143 for young vs old and mutant vs wildtype comparisons; see below), does not suffer from this potential issue. However, the choice of RNA-sequencing analysis platform can significantly influence the detection of differentially expressed genes (DEGs) when analysing large datasets (*133*). Specifically, a recent study evaluating the performance of the two most popular tools, DESeq2 (*134*) and edgeR (*135*), in identifying DEGs in large-sample studies found that the DEGs they detected were anticonservative (*136*). This observation prompted us to use a cautious approach when classifying genes as DE, analysing our data using both EdgeR (v4.6.2) and DeSeq2 (v1.48.1). For both platforms, we used the design matrix: ∼ *Age* ∗ *Deletion. Status*, where Age and Deletion.Status are factors with two levels each (Old (O) and Young (Y), and Wildtype (wt) and Mutant (mu). Additive effects of age and deletion status were extracted using the contrasts ((O_wt + O_mu) − (Y_wt + Y_mu))/2, and ((*Y*_*mu* + *O*_*mu*) − (*Y*_*wt* + *O*_*wt*))/2, respectively. We also applied a more stringent false discovery rate (FDR) cut-off (0.01) than the cut-off (0.05) that is commonly used. For Deseq2, differential expression analysis was performed using the DESeq function, followed by estimation of log_2_ fold-change using the *lfcshrink* function with the adaptive shrinkage estimator from the *ashr* package (*137*). In edgeR, the analysis was performed by first filtering low-count genes with *filterByExpr*, then fitting the design with glmQLFit. Significant DEGs were extracted using the function *glmQLFTest*. While we found a high degree of overlap in the DEGs detected by the two packages (Figure S4), indicating robustness, we took a more conservative approach and retained only the DEGs detected by both packages for further analyses. The effects of age and deletion on gene expression (log_2_ fold-change) were extracted from the DESeq2 platform, given the robustness of estimates provided by *lfcshrink* (*137*, *138*).

Gene Ontology (GO) term overrepresentation analyses were performed separately for up- and downregulated DEGs for both age and condition using the *enrichGO* function from clusterProfiler (v4.16.0) (*139*). Following the GO analysis, the DEGs were classified into two groups: genes that belonged to at least one significant GO term were classified as “enriched” and those that did not were classified as “unenriched”, resulting in four categories for downstream analyses - upregulated-enriched, downregulated-enriched, upregulated-unenriched, downregulated-unenriched. We predicted that the enriched genes would be differentially expressed as a plastic response to a decline in condition and/or heightened stress and would therefore exhibit distinct characteristics compared with unenriched genes.

The distribution of “null” expectations of overlapping GO terms between age and condition was simulated for up- and downregulated genes using a Monte Carlo permutation (N=15000). For each simulation, the same number of genes up- or downregulated by condition (200 or 293, respectively) were sampled from the genes up- (or down-) regulated by age, followed by GO analysis of the gene set. The number of overlapping GO categories between the sample set and age DEGs was then calculated to establish the null distribution, and an exact P-value ([N_permuted overlap > observed overlap_ + 1]/[N + 1]) was calculated from it.

A similar simulation approach was used to test for directional clustering of condition effects. We first analysed GO categories enriched for age-only DEGs. For each category, we used a one-sided binomial test (P < 0.05) to determine whether genes not significantly affected by condition showed a logFC direction matching the significant age effect more often than chance would predict. Null expectations were established by permuting the direction of condition logFCs for genes that belong to an age-related GO category but are not significantly affected by condition, 15000 times for both downregulated and upregulated categories.

Tests for gene set overlap between DEGs due to age and condition were performed in two steps. We first tested whether the 414 DEGs that were significantly overlapping between age and condition (at the initial statistical threshold mentioned above) were more than expected by chance. This analysis was further refined by cumulatively binning the significant genes into 50 equidistant bins of increasing stringency based on FDR-adjusted P values, dividing the genes into up- and downregulated sets, and performing enrichment tests for each bin. All enrichment tests were performed using a one-tailed hypergeometric test.

Similarly, the effect size (log_2_Fold-change) correlations were initially performed on the 414 significant DEGs, followed by refinement using an adjusted P-value threshold; in this case, the number of bins was reduced to 10 to ensure enough genes in each bin for meaningful correlation tests. Additionally, we tested effect-size correlations between age and condition for enriched and unenriched age-related DEGs that were not significantly affected by condition. Here also, the genes were divided into 10 bins with increasing statistical stringency. All correlation tests were performed using Spearman’s rank correlation test as implemented by the function *cor.test*.

### Analysis for Association with Histone Modifications

To assess whether differentially expressed genes (DEGs) were associated with specific chromatin states, we utilized a recently published histone modification data set from young *D. melanogaster* female head tissue (*55*). From this source, we extracted the binary classification (present or absent) for the activating mark H3K4me3 and the repressive mark H3K27me2/me3 for all genes included in our analysis.

We then determined the proportion of genes associated with each histone mark within distinct sets of up- and downregulated DEGs. For age-related DEGs, we used chi-squared tests to compare these proportions between functionally enriched and unenriched gene sets. A similar approach was taken for condition-related DEGs, where we first analyzed the complete set of DEGs and subsequently performed a refined analysis focusing only on the subset of functionally enriched genes. In all cases, the proportions for each gene set were statistically compared against a baseline composed of genes not significantly affected by the respective treatment (age or condition).

### Analyses of genes with tissue-biased expression

Tissue-specific normalized expression (Fragments Per Kilobase of transcript per Million mapped reads; FPKM) values were obtained from the FlyAtlas2 RNA-Seq database (*67*) for 11 female tissues (head, ovary, mated spermatheca, hindgut, midgut, crop, salivary gland, Malpighian tubule, virgin spermatheca, fat body and heart). Brain and eye were excluded to avoid redundancy. Using this data, z-scores for female head were calculated as a metric for head-biased gene expression, i.e., increasing values (range = [-1.16, 3.02]; median = 0.98) indicate increased gene expression bias towards head. We next classified genes with z-score >1.5 as head-biased and z-score <-0.5 reverse-head-biased. A smaller value for classifying genes as reverse-head-biased was selected due to the paucity of the reverse-head-biased genes in our data for obvious reasons (they were collected from head samples), and we made the choice to keep the thresholds roughly equidistant from the median. Following the binary classification, two-sample proportion tests were performed using the function *prop.test* for testing the difference in their proportional representation between up- vs downregulated by age and condition respectively.

To test if increased tissue bias led to proportional downregulation and if such an effect depended on whether the genes belonged to an enriched or unenriched class, we employed bootstrap regression. We defined a custom bootstrap function to sample the data with replacement and fit linear models using the following function y ∼ Enrichment * z-score, where y represents the log_2_ fold change in expression due to either age or condition, and Enrichment (factor) represents whether the genes belonged to the enriched or unenriched category for aging DEGs and dummy coded to reflect if they included all genes or only enriched genes for condition DEGs. We set up a parallel backend using the *doParallel* package (https://github.com/RevolutionAnalytics/doparallel) and ran the bootstrap function in parallel for 5000 iterations. We extracted the regression coefficient from each bootstrap (b_bootstrap_) and used the 97.5% (upper) and 2.5% (lower) quantiles as confidence intervals (CI) to test whether b_bootstrap_ estimates were significantly different from zero for each group.

### Analyses of genes with sex-biased expression

The identity of sex-biased genes can vary substantially among populations within the same species (*41*). Thus, to ensure comparability across studies, we referenced the dataset from Malacrinò et al. (2022) (*41*), which profiled sex-biased gene expression in 5-day-old females from the wild-type Dahomey population, the same (young) age and genetic background used in the present study and detected 540 female-biased genes. Enrichment of these genes in our age- and condition-induced enriched and unenriched DEGs were tested using chi-squared and hypergeometric tests, as appropriate.

### Association between aging-related gene expression and coefficients of variation (CV) in gene expression

To assess variability in gene expression across samples, we calculated the coefficient of variation (CV) for each gene using normalized expression values. Raw counts were first normalized using the cpm function from the edgeR package (v4.6.2) after adding a pseudocount of 1 and then log2 transformed. Genes were filtered to include those with significant age-related expression changes. For each gene, we computed the mean and standard deviation of CPM values across samples, stratified by sample age (old, young). The CV was calculated as the ratio of standard deviation to mean CPM. Statistical differences in CV were tested using a linear model with the formula *log (CV) ∼ Enrichment*Effect*SampleAge* where *Enrichment, Effect* and *SampleAge* represent GO enrichment category (enriched, unenriched), age-related differential expression status (upregulated, downregulated), and sample age (old, young) as interacting predictors, respectively. Type III ANOVA was performed using the car package (*140*), and estimated marginal means were extracted using the emmeans package (*141*). Pairwise contrasts were adjusted using the Bonferroni method.

### Association between different classes of genes and evolutionary rates

To investigate whether genes differentially expressed with age or condition exhibit distinct evolutionary rates, we analysed the relationship between gene expression categories and the ratio of nonsynonymous to synonymous substitutions (dN/dS), for which the data were obtained from (*88*). Briefly, orthologs from 15 species in the melanogaster subgroup were obtained using a local installation of OrthoDB (*142*), followed by protein-based alignments and backtranslation using MAFFT. The resulting orthology relationships formed the basis for producing protein-based alignments using the MAFFT L-INS-I algorithm (*143*). dN and dS values for each ortholog were estimated using PAML (*144*).

Because gene expression level and tissue specificity correlate with dN/dS values, we used tissue specificity value (τ) and female whole-body expression and their interaction as covariates. These values were obtained from FlyAtlas2. For age-related DE the gene set was categorized based on effect and GO enrichment classification, resulting in composite categories (i.e., Downregulated Enriched, Upregulated Enriched, Downregulated Unenriched and Upregulated Unenriched). For condition-related DE, the gene set was first categorized based on effect (i.e., Up- or Downregulated), followed by a secondary analysis where they were refined to only enriched genes. Genes not significantly affected by age or condition were labelled as “NS”. We modelled dN/dS using Bayesian regression implemented in the brms package (v2.21.0)(*145*). Separate models were fitted for age- and condition-associated genes. In both models, dN/dS was the response variable, and the predictors included gene category, tissue specificity (τ), and whole-body expression level, with an interaction term between τ and expression. A hurdle lognormal distribution was used to account for the zero-inflated and skewed nature of dN/dS. Models were run with 5 chains, 15,000 iterations per chain. Parallel computation was enabled using the future (*146*) and parallel (*147*) packages.

Unless otherwise specified, all analyses were performed in R (v 4.5) (*147*). All figures were made using packages ggplot2 (*148*), ggpubr (*149*) and ggrepel (*150*).

## Data availability

All RNA-Seq data including raw-reads will be made available at the European Nucleotide Archive ENA data repository. All the analysis scripts are available at: https://github.com/nahseezila/Aging-deletion

## Acknowledgements

We thank Miruna Dumea for excellent assistance with both backcrossing of deletions into our lab population and rearing of the flies used in the experiment. The authors acknowledge support from the National Genomics Infrastructure in Stockholm funded by Science for Life Laboratory, the Knut and Alice Wallenberg Foundation and the Swedish Research Council. The computations and data handling were enabled by resources in project [NAISS 2023/23-42; NAISS 2023/22-93; NAISS 2025/5-81; NAISS 2025/6-168] provided by the National Academic Infrastructure for Supercomputing in Sweden (NAISS) at UPPMAX and PDC, funded by the Swedish Research Council through grant agreement no. 2022-06725. Stocks obtained from the Bloomington Drosophila Stock Center (NIH P40OD018537) were used in this study.

