## Supplementary file 1 for "The aging transcriptome: a condition-dependent response to a deteriorating soma"

**Contents:**

1. Additional methods
2. Supplementary figures S1, S2, S3
1. **Additional methods: Filtration criteria of genes used in the study:**

The deletions used in this study present several artifacts that could explain differences in the expression of a subset of genes. To avoid such confounds when interpreting changes in gene expression, the following criteria were used to remove genes from the dataset prior to final differential expression and downstream analyses:

First, we removed the 554 genes (Table S1) included in one of the 18 deletions, as most of these distort gene expression between deletion and wild type samples simply due to differences in copy number.

Second, we removed the gene *white* (*w*) from our analyses, as flies carrying a deletion were made phenotypically different from wild type flies by having the deletion flanked with *mini-white* (a functional copy of *w*), on an otherwise shared genetic background that was homozygous for a non-functional version of *w*. Flies carrying a deletion hence had three copies of this gene, potentially causing gene expression differences simply due to differences in copy number. Overexpression of *w* could also affect genes that physically interact with *w*. Consequently, we identified and removed 117 known primary interactors of *w* from the STRING database (Szklarczyk et al., 2023).

Third, following an initial differential expression analysis, comparing wild type to deletion flies, using DESeq2 and subsequent gene ontology enrichment analysis (see Methods for GO analysis details), we identified 18 genes enriched in several categories related to vision-related processes (Figure SA). These genes are downregulated by age but upregulated in the presence of deletions (Figure SB). A plausible explanation for the observed expression changes is the presence of the extra copy of the *w* gene white. To test this hypothesis, we used R function “prcomp” (R Core Team, 2022) to perform a principal component analysis (PCA) of the standardized counts per million (CPM) values calculated by the function cpm in package edgeR (Chen et al., 2025) of these genes. The PCA revealed that the first principal component (PC1) explained most of the variation (85.7%; Figure SC), indicating substantial co-expression. Regression analysis of PC1 scores against expression of *white* in wild type samples, after controlling for the effect of age, showed a highly significant positive association (Figure SD), suggesting that the expression of these genes is related to *white* expression and therefore also affected by the presence of the *mini-white* marker in flies carrying a deletion. Consequently, we therefore also removed these 18 genes from the analyses.

Finally, we removed genes with read count less than 2 in more than 50% of the samples

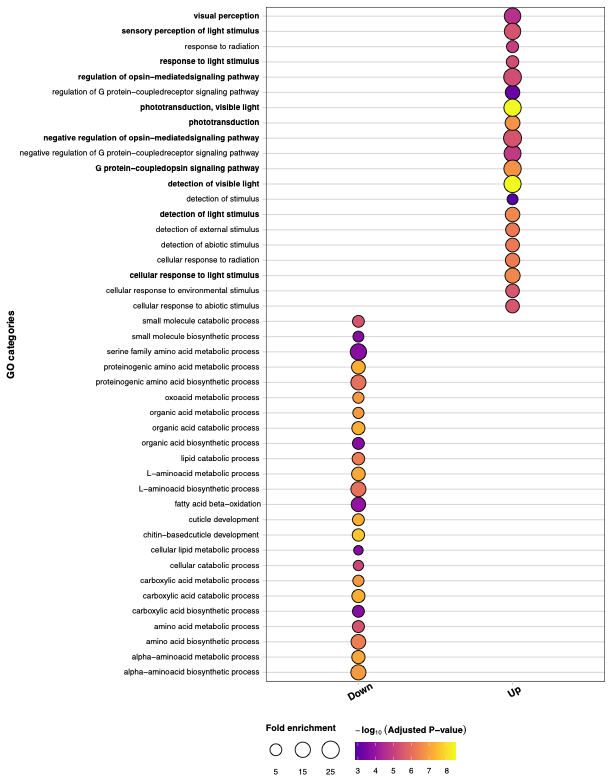

**Figure SA:** Top 20 gene ontology (GO) categories for genes differentially expressed by condition prior to filtration. The x-axis represents the direction of change. The y-axis represents GO: biological processes (BP). The boldface GO: BP categories relate to photosensitivity, and genes belonging to these categories were removed. The heat of the color corresponds to significance level, and the sizes of the circles correspond to fold-enrichment.

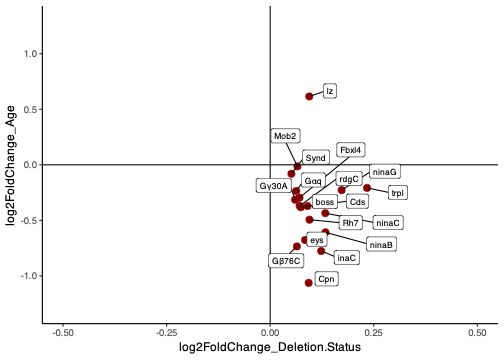

**Figure SB:** Effect (log_2_ fold-change) of age (y-axis) and deletion (x-axis) on the 18 photosensitivity related genes. All but one of these genes is discordantly affected by age and condition, i.e., down by age, and upregulated by condition.

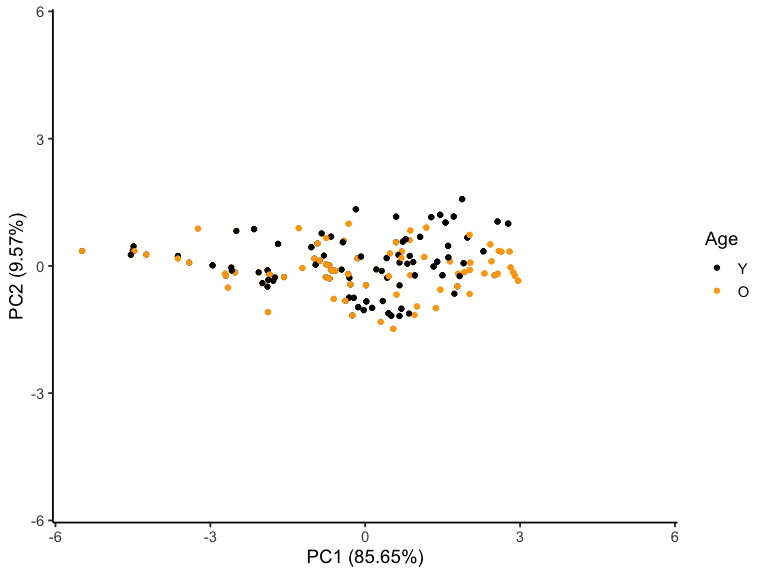

**Figure SC:** Principal Component Analysis (PCA) of counts per million values of the 18 photosensitivity related genes in wildtype samples showing separation along PC1 (85.65%) and PC2 (9.57%), with samples grouped by age category (Y = Young, O = Old).

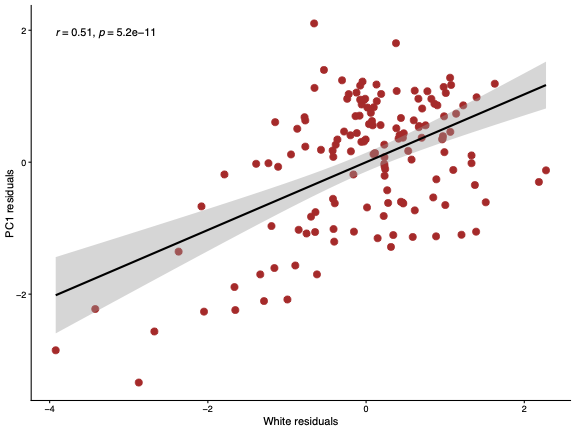

**Figure SD:** Residual regression plot showing the relationship between PC1 residuals and *white* residuals. The residuals were calculated from a regression over age to account for any effect of age on their gene expression.

2. **Supplementary figures**

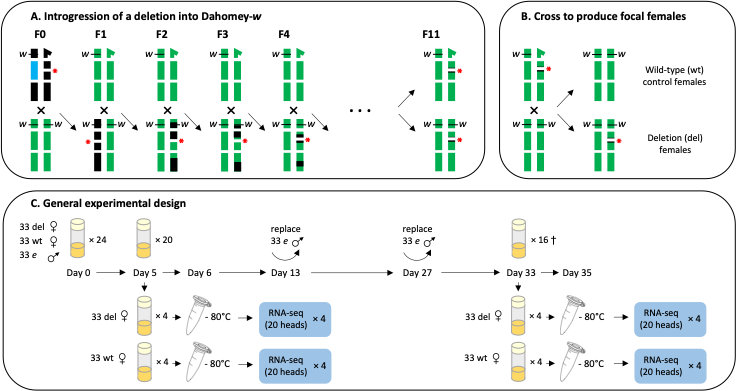

**Supplementary figure 1.** Overview of experimental procedures. **(A)** Crosses used to introgress one deletion into Dahomey-*w*, a copy of the long-term laboratory adapted population Dahomey homozygous for the recessive eye-color marker *white* (*w*). Top row: male genotypes; bottom row: female genotypes. Sex chromosomes are shown at the top, major autosomes (2 and 3) below. The 4^th^ “dot” chromosome is omitted for brevity. Males carrying the deletion (depicted with a star and as a gap on chromosome 2) balanced over a balancer chromosome (blue) were crossed to virgin Dahomey-w females. Virgin daughters carrying the deletion were subsequently crossed to Dahomey-w males. The deletion is flanked by mini-white [w^+mC^] (red star), rescuing the red-eyed phenotype and allowing identification of deletion carriers. This scheme was repeated for all 18 deletions for ≥10 generations, replacing the original deletion background (black) with Dahomey (green). **(B)** Cross to generate focal deletion and wild-type females used in the experiment. **(C)** Experimental setup: 33 focal wild-type and 33 deletion females were housed with 33 ebony (e) males in each of 24 vials. At day 5, 33 focal females per type (from 4 vials) were sorted under light CO₂ anaesthesia and kept separately for 1 day. At day 6, they were flash-frozen and stored at -80°C. Remaining females were aged to day 35, with ebony males replaced twice; sorting was repeated at day 33 for 4 new vials. Heads from 20 females per sample were pooled for RNA-seq.

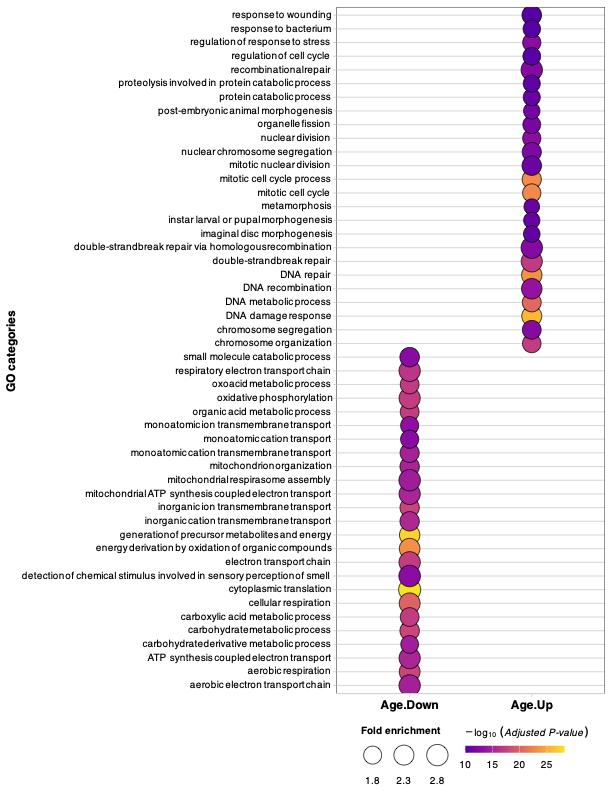

**Supplementary figure 2.** Enriched gene ontology (GO: biological process) categories for DEGs in young and old flies. The x-axis represents the gene-classes that are down- and upregulated in old flies. The y-axis represents top 20 GO: BP categories for each set of genes. Consistent with previous studies, the downregulated genes are enriched for metabolic processes, mitochondrial function, and cytoplasmic translation, whereas the upregulated genes are enriched for repair and stress response related process. The color gradient represents significance levels (adjusted *P-*values). The size of the dots represents degree of enrichment (log_2_-fold) above random expectation.

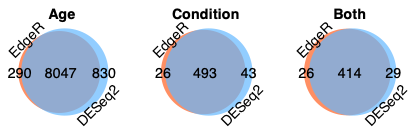
**Supplementary figure 3.** Venn diagrams showing high degree of agreement in detecting differentially expressed genes (DEGs) using two different platforms: EdgeR (orange) and DeSeq2 (light blue). All three DEG-sets detected for age, condition, and age-condition overlap show considerable intersection (dark blue) between the two platforms. The numbers represent the number of DEGs either in the intersection (middle) or belonging uniquely to the corresponding platforms.
